# Structural (dys)connectivity associates with cholinergic cell density of the nucleus basalis of Meynert in Alzheimer’s disease

**DOI:** 10.1101/2021.08.02.454716

**Authors:** Chen-Pei Lin, Irene Frigerio, Baayla DC Boon, Zihan Zhou, Annemieke JM Rozemuller, Femke Bouwman, Menno M Schoonheim, Wilma DJ van de Berg, Laura Jonkman

## Abstract

Cognitive deficits in Alzheimer’s disease, specifically amnestic (memory dominant) deficits, are associated with cholinergic degeneration in the basal forebrain. The cholinergic nucleus within the basal forebrain, the nucleus basalis of Meynert, exhibits local atrophy and reduced cortical tract integrity on MRI, and reveals amyloid-β and phosphorylated-tau pathology at autopsy. To better understand the pathophysiology of nucleus basalis of Meynert atrophy and its neocortical projections in Alzheimer’s disease, we utilized a combined post-mortem *in-situ* MRI and histopathology approach. A total of 19 Alzheimer’s disease (10 amnestic and 9 non-amnestic) and 9 non-neurological control donors underwent 3T T1-weighted MRI for anatomical delineation and volume assessment of the nucleus basalis of Meynert, and diffusion-weighted imaging for microstructural assessment of the nucleus and its projections. At subsequent brain autopsy, tissue dissection and immunohistochemistry were performed for amyloid-β, phosphorylated-tau and choline acetyltransferase. Compared to controls, we observed an MRI-derived volume reduction and altered microstructural integrity of the nucleus basalis of Meynert in Alzheimer’s disease donors. Furthermore, decreased cholinergic cell density was associated with reduced integrity of the nucleus and its tracts to the temporal lobe, specifically to the temporal pole of the superior temporal gyrus, and the parahippocampal gyrus. The association between cholinergic cell density and alteration to cortical tracts was specific for amnestic, compared to non-amnestic Alzheimer’s disease donors. Our study illustrates that the nucleus basalis of Meynert is severely affected in amnestic Alzheimer’s disease, both in terms of pathology within the nucleus, but also in terms of damage to its cortical projections, specifically to the temporal lobe, which may contribute to the observed cognitive deterioration.

## Introduction

Alzheimer’s disease is a heterogeneous disease characterized by memory deficits and cognitive decline with specific patterns of neurodegeneration. Clinically, a distinction can be made between an amnestic and non-amnestic subtype. In amnestic Alzheimer’s disease, memory deficits are among the first symptoms, and in non-amnestic Alzheimer’s disease, memory is initially spared, but symptoms of visuospatial impairment, aphasia, or behavioural/dysexecutive dysfunction are more prominent^1^. Pathologically, Alzheimer’s disease is characterized by abnormal aggregation of amyloid-beta (Aβ) forming amyloid plaques, and phosphorylated-tau (p-tau) forming neurofibrillary tangles (NFT)^2,3^. Furthermore, one of the earliest sites of neurodegeneration is within the cholinergic nucleus basalis of Meynert (NbM), which is located in the substantia innominata of the basal forebrain^4^. Aβ and NFT accumulation has been associated with cholinergic cell loss in the NbM, which in turn associated with decreased cortical acetylcholine and plays an important role in disease-related cognitive decline, specifically in memory and attention processing^5–11^. Neuroimaging of the NbM has shown to be useful, for instance NbM volume can predict cognitive improvement after intake of galantamine, a common drug treatment targeting the cholinergic system in Alzheimer’s disease^12^. Moreover, deep brain stimulation of the NbM is explored as a possible treatment option in Alzheimer’s disease, by restoring the cholinergic transmission that is crucial in mediating memory processing^13–15^. It is therefore important to better understand the interplay between NbM protein aggregation, cholinergic cell loss, neuroimaging markers, and cognition in Alzheimer’s disease. This may help in evaluating and monitoring disease-related alterations, as well as therapeutic effects of cholinergic-targeting interventions.

*In vivo* MRI-measured volume of the NbM is thought to be sensitive to cholinergic degeneration, as illustrated by the decreased volume of the NbM in Alzheimer’s disease patients compared to controls^11,16^. Moreover, recent advances in diffusion MRI measurements, specifically of fractional anisotropy (FA) and mean diffusivity (MD), allows us to not only examine the microstructural integrity within the NbM, but also the microstructural integrity of its cholinergic projections^17^. Cholinergic projections to the frontal and temporal lobe have shown to be predominantly affected, and associate with cognitive dysfunctions^18–21^. Nevertheless, in these studies, the pathological substrate underlying these MRI-measured alterations of the NbM and their cortical projections, were not explicitly addressed.

To this end, the current study investigated the association between histopathological measures of cholinergic cell density, Aβ and p-tau, with within-subject post-mortem MRI-derived NbM volume, microstructural integrity, and cortical projections, in Alzheimer’s disease and non-neurological control donors. We hypothesized that, aside from the volume and microstructural integrity in the NbM, its cortical projections specifically to the frontal and temporal cortex are affected and associate with pathological measures. In addition, due to the clinical and pathological heterogeneity in Alzheimer’s disease, we explored the MRI and histopathological associations in clinically defined amnestic and non-amnestic subtypes, and hypothesized that the amnestic subtype is especially affected due to the cholinergic involvement in memory processing. The current study may lead to a better understanding of the pathophysiology underlying NbM atrophy and projections in *in vivo* imaging.

## Materials and methods

### Donor inclusion

A total of 28 brain donors, 19 Alzheimer’s disease and 9 non-neurological control donors, were included in the study, with written informed consent for the use of their brain tissue and medical records for research purposes. Alzheimer’s disease donors were included in collaboration with the Alzheimer center Amsterdam and the Netherlands Brain Bank (NBB; http://brainbank.nl). During life, patients were screened according to the screening protocol of the Amsterdam Dementia Cohort^22^. Diagnosis was made in a multidisciplinary team according to NINCDS-ADRDA criteria^23^. An Alzheimer’s disease subtype distinction could be made between amnestic (N=10) patients, and non-amnestic (N=9) patients according to the IWG-2 criteria^24^, with the latter one being subdivided into 6 patients clinically diagnosed with the behavioural/dysexecutive and 3 patients with posterior cortical atrophy^24,25^. Non-neurological controls were included at the department of Anatomy and Neurosciences, Amsterdam UMC, location VUmc, following the Normal Aging Brain Collection Amsterdam (NABCA; http://nabca.eu) pipeline^26^. Neuropathological diagnosis of all donors was confirmed by an expert neuropathologist (A.R.) and performed according to the international guidelines of the Brain Net Europe II (BNE) consortium (http://www.brainnet-europe.org). All donors underwent post-mortem *in-situ* MRI and brain autopsy. The study design is summarized in Fig.1.

**Figure 1.**
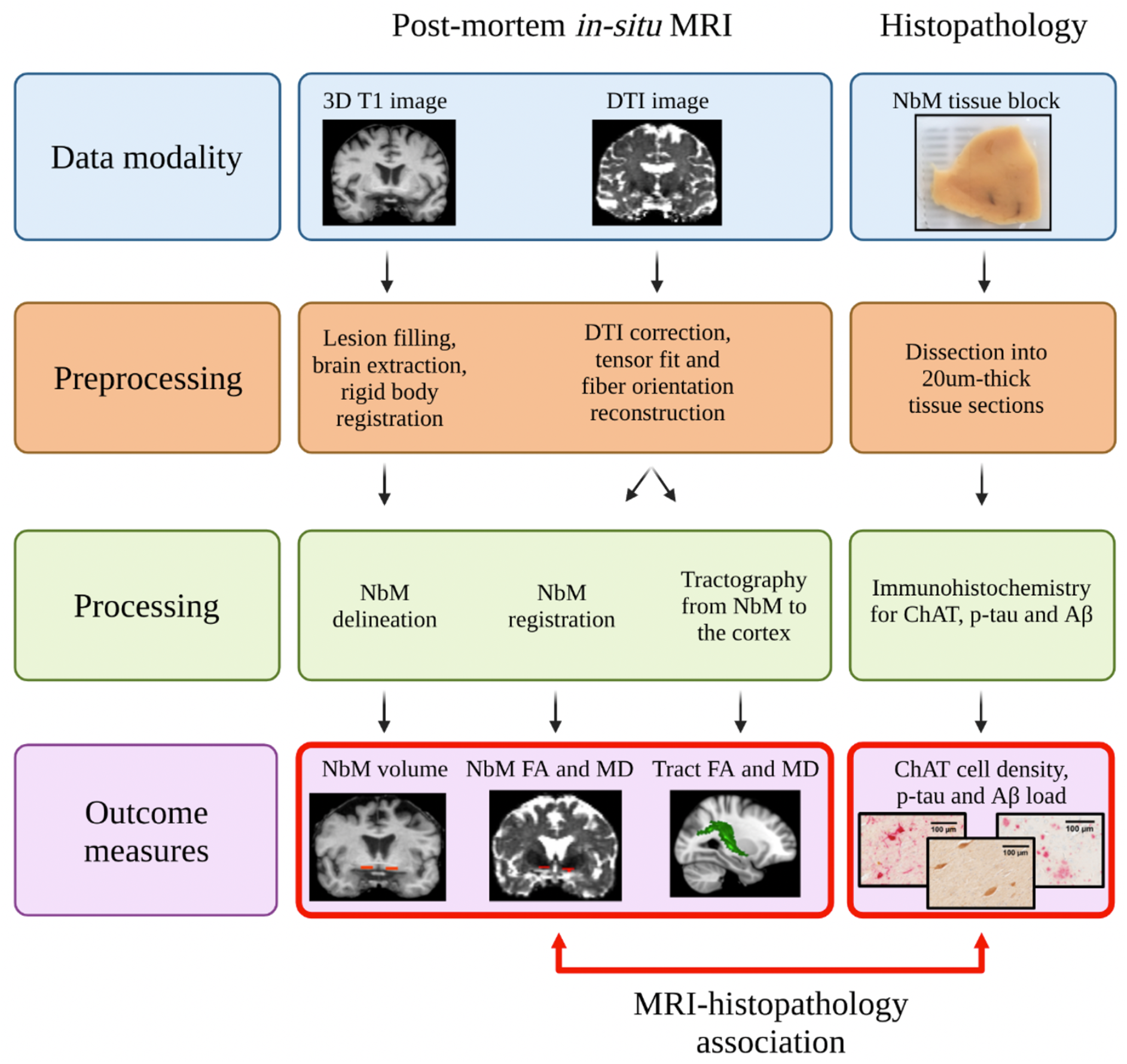
Study design of combined post-mortem *in-situ* MRI and histopathology. The red dotted arrow indicates the statistical analysis between MRI and histopathology outcome measures. Abbreviations: DTI=diffusion tensor images, NbM=nucleus basalis of Meynert, FA=fractional anisotropy, MD=mean diffusivity, ChAT=choline acetyltransferase, p-tau=phosphorylated tau, Aβ=amyloid-beta.

### MRI acquisition

Post-mortem MRI data was acquired by whole-brain *in-situ* (brain still in cranium) scanning on a whole-body 3T MR scanner (Signa-MR750, General Electric Medical Systems, United States) with an eight-channel phased-array head-coil^26^. Details of the MRI protocol with 3D-T1-weighted images, fluid attenuated inversion recovery and diffusion tensor images (DTI) sequences can be found in Supplementary Materials.

### MRI analysis

#### Structural image processing

To minimize the impact of age-related white matter abnormalities (e.g., vascular change) on automated segmentations, the 3D T1 images were lesion-filled^27^, as previously described^28^. Subsequently, normalized brain volumes of the whole brain, white matter and gray matter were estimated from 3D T1 images using SIENAX^29^, FMRIB Software Library (FSL) tools version 5.0.9 (https://fsl.fmrib.ox.ac.uk/fsl/). In addition, each hemisphere was parcellated into 39 anatomical regions using the automated anatomical labeling (AAL) atlas^30^, and transformed from 3D T1 to diffusion space using Boundary-Based registration (BBR) and FMRIB’s Integrated Registration and Segmentation Tool (FLIRT) for further diffusion analysis.

#### NbM delineation on MRI

3D T1 images were registered to MNI orientation using an affine rigid-body transformation. In this standardized orientation, the NbM was delineated (both left and right hemisphere) in each subject, using five consecutive coronal sections of 1mm, based on the method described previously^31–33^. In brief, the first delineating section was identified at the level of crossing anterior commissure. The dorsal border of the NbM aligned with the most ventral aspect of the globus pallidus, while the ventral border of the NbM was the final row of voxels before the cerebrospinal fluid. The lateral border of NbM was delineated along the medial aspect of the putamen, whereas the medial border was demarcated following the extended ventrolateral outline of the internal capsule to the base of the brain (Fig.2A and Fig.2B). Left and right NbM volumes were calculated and normalized for head size using the V-scaling factor from SIENAX. Subsequently, the NbM was reversely transformed to native 3D T1 space, and co-registered to diffusion space using BBR parameters.

**Figure 2.**
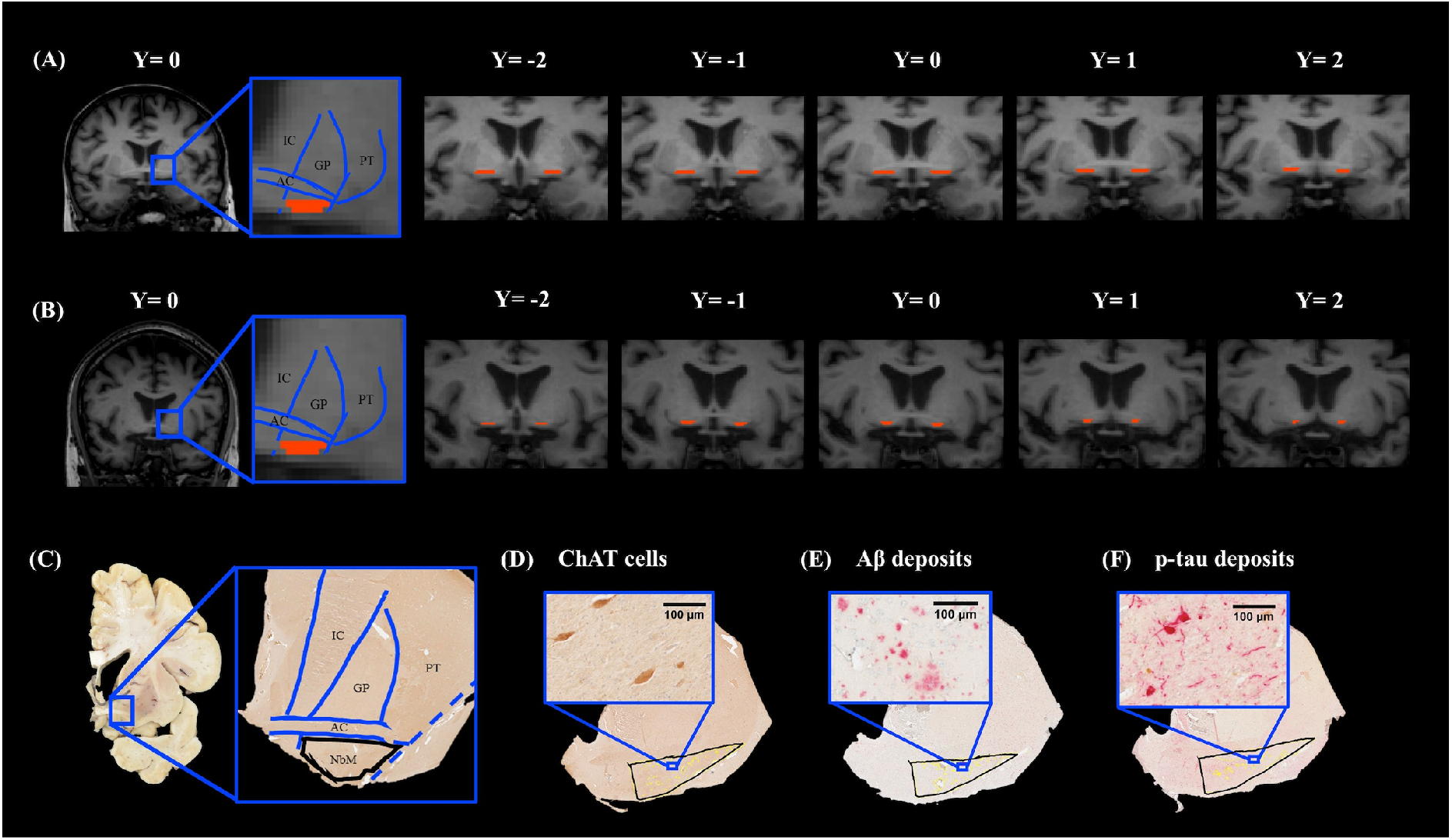
NbM delineation in MRI T1 coronal scans and histological sections. Illustration of NbM delineation in five consecutive T1 coronal images of a control (A) and an AD donor (B). On the image with the highest signal intensity of visible anterior commissure (AC), denoted as Y=0, the NbM was delineated according to the standard guideline based on anatomical landmarks of the AC, internal capsule (IC), globus pallidus (GP), putamen (PT). Subsequently, the NbM was delineated on two adjacent images forwards and backwards (Y=-2 to Y=2). (C) Illustration of NbM delineation in histological sections of a control. Delineation was performed primarily on ChAT single stained sections, with illustrated standard guidelines based on the same anatomical landmarks as in (A). In an Alzheimer’s disease donor, the delineated NbM exhibits ChAT immunoreactivity with visible ChAT cells in (D), Aβ deposit in (E) and p-tau deposit in (F) with visible p-tau tangles and threads. Abbreviations: ChAT=choline acetyltransferase, p-tau=phosphorylated tau, Aβ=amyloid-beta.

#### DTI processing

DTI were first corrected for eddy current induced geometric distortion^34^ and fitted for diffusion tensors^35^. Subsequently, diffusion orientation distributions were modeled using FDT (part of FSL 5.0.9). The delineated NbM ROI was overlayed onto the FA and MD diffusion maps to derive the FA and MD values of the NbM^36^. Tracts between the NbM and cortical atlas regions were determined using probabilistic tractography. For this, BEDPOSTX, a function from FDT, was used to model the distribution of fiber orientations at each voxel yielding the voxel-wise diffusion orientations for performing probabilistic tractography.

#### Probabilistic tractography

Probabilistic tractography was performed using ProbTrackX2 (FDT, FSL 5.0.9) with default settings and 5000 sampling fibers. The cortical tracts were reconstructed in each hemisphere separately with left or right NbM as seed regions of interest (ROIs). Subsequently, tracts were binarized and FA and MD of tracts were calculated by overlaying the tracts onto the diffusion maps. In a stepwise fashion, to limit the number of comparisons, we first addressed tracts from the NbM to the cingulum and cortical lobes (frontal, temporal, parietal, occipital cortex and insula)^37^, see Supplementary Table 1 for a list of composit AAL regions. For tracts that showed significant associations between FA or MD and pathological measures, we subsequently addressed the integrity of tracts to the subregions of these cortical lobes (e.g., superior, middle and inferior temporal gyrus; Supplementary Table 1).

### NbM tissue sampling, processing and quantification

After post-mortem *in-situ* MRI, the donors were transported to the mortuary for brain autopsy. After the brain was extracted from cranium, the right hemisphere was fixed in 4% formalin for four weeks and subsequently dissected for specific anatomical regions according to a standardized protocol^26,38^. Tissue blocks were retrospectively collected from each case. Based on the Dickson sampling scheme^39^, tissue blocks with visible substantia innominata that contains the NbM was sampled in a corona plan where the substantia innominata was visible underneath the anterior commissure at the level of the caudate-putamen. The blocks were subsequently paraffin embedded, followed by immunohistochemistry.

#### Immunohistochemistry for cholinergic neurons, Aβ and p-tau

For detailed methods, see Supplementary Materials. In brief, paraffin-embedded tissue NbM blocks were cut at 20 μm for 30 consecutive sections and stained for single Choline Acetyltransferase (ChAT) and double ChAT/Aβ (6F/3D) and ChAT/p-tau (AT8). From the first section with visible anterior commissure and substantia innominata, three sections with a distance of 200 um in between were included per staining. The primary antibodies used for each staining are shown in Supplementary Table 2. Finally, ChAT was visualized with 3,3’-Diaminovenzidine (DAB, Dako) imidazole, whereas p-tau (AT8) and Aβ (6F/3D) were visualized using liquid permanent red followed by counter-staining with haematoxylin, and mounted with Entellan.

#### NbM delineation on tissue sections

Tissue sections of four Alzheimer’s disease cases had to be excluded due to incorrect anatomical orientation or tissue disintegration during the staining process. Immunostained sections were digitally scanned with the Vectra Polaris Quantitative Pathology Imaging System (PerkinElmer, USA) at 20x magnification. The NbM was manually delineated on the ChAT-stained sections using Fiji ImageJ Version 1.52r (https://imagej.nih.gov/ij), which was subsequently used as template for delineation in the ChAT/p-tau and ChAT/Aβ double stained sections (Fig.2C-F). Based on previous literature^4,40,41^, the ROI for NbM was defined independent of the cholinergic cells, but according to the anatomical landmarks of neighboring structures so that the disease-related cholinergic loss would not affect the selected area. Similar to the MRI delineation, the ventral aspect of the anterior commissure serves as the dorsal border, whereas the base of the brain serves as the ventral border of the ROI. The lateral border was delineated following the ventrolateral outline of the global pallidus that intersects the anterior commissure and further extends to the base of the brain, while the medial border aligns the extended lateral outline of internal capsule to the base of the brain. To account for the heterogeneous distribution of cholinergic cells within NbM, the sections were further subdivided into “pre-anterior”, “anteroromediate” and “antero-intermediate” NbM based on the emergence of anterior commissure and neighboring anatomical structures^4,7,41–43^ and visualized in Supplementary Fig 1. The sections were denoted as “pre-anterior NbM” if the ChAT cells were underneath the visible and continuous anterior commissure that is ventral to the globus pallidus and rostral to decussation level. The sections were denoted as “anteroromediate NbM” if the ChAT cells were underneath or adjacent to the elongated and continuous anterior commissure. The sections were denoted as “antero-intermediate NbM”, if the ChAT cells were underneath or adjacent to the tip of the rostral anterior commissure and underneath the putamen. As these subsectors were unevenly distributed in the Alzheimer’s disease and control group, this distinction is included as a covariate in the statistical analysis.

#### Quantification of cholinergic cell density and pathological load

After ROI delineation, ChAT, Aβ and p-tau were quantified using in-house Image-J scripts. For ChAT cell density, circular objects with a size between 20-200 μm in diameter were selected and ChAT cell count per mm^2^ was calculated. For Aβ and the p-tau, the load (%area) showing immunopositive signal within the ROI was calculated (Supplementary Fig.2). All derived measures were averaged across the three sections for each immunostaining. In summary, our immunohistochemical outcome measures after averaging across three sections were ChAT cell density, Aβ load, and p-tau load.

### Statistical analysis

Statistical analysis was performed using IBM SPSS 22.0 for Windows (SPSS, Inc., Chicago, IL, USA). All the statistical variables were tested for normality. Non-normally distributed data was log transformed, including FA, MD, ChAT cell density, Aβ and p-tau load. Chi-square test was used to investigate group difference between Alzheimer’s disease donors and non-neurological controls, and between Alzheimer’s disease subtypes (amnestic and non-amnestic) for categorical variables, e.g., gender and pathological staging. General linear models (GLM) were used for the aforementioned group differences in MRI-derived outcome measures (NbM volume, FA and MD, as well as FA and MD of tracts to the cortex), and histopathological outcome measures (ChAT cell density, Aβ and p-tau load). We applied partial correlations and linear mixed models to examine the associations between the above mentioned MRI and histopathology-derived outcome measures within the right hemisphere. Age, gender and postmortem delay (PMD) were included as covariates in all statistical analysis^44^. The NbM subsectors, as mentioned in previous paragraph, were included as covariate in statistical analysis of ChAT cell density. NbM volume was included as covariate in statistical analysis of NbM tracts^45^. All data were corrected for multiple comparisons with Bonferroni correction and false discovery rate (FDR)^46^.

## Results

### Study cohort

Clinical, neuropathological and radiological characteristics of each group are shown in Table 1, and details of each donor is shown in Supplementary Table 3. There were no significant differences in age and PMD between Alzheimer’s disease donors and controls, however, the Alzheimer’s disease group had more males than the control group (p=.02). Normalized total brain volume and normalized white matter volume were comparable between groups. As expected, the Alzheimer’s disease donors had lower normalized grey matter volume (p=.046), higher Braak NFT stages (p<.001), Thal phases (p<.001) and ABC scores (p<.001) than controls, while APOE genotype did not differ between groups (p=.453).

**Table 1.** Clinical, radiological and pathological characteristics of donors.

|  | **Controls** | **AD** |
| --- | --- | --- |
| **Clinical characteristics** |  |  |
| **N** | **9** | **19** |
| **Gender F/M (%M)** | 4/5 (56%) | 4/15 (80%) |
| **Age at death**<br>years, mean ± SD | 70.8 ± 8.8 | 67.4 ± 11.6 |
| **Disease duration**<br>years, mean ± SD | - | 8.1 ± 4.8 |
| **PMD, mean (hr:min) ± SD (hr)** | 9:10 ± 3 | 7:23 ± 2 |
| **Radiological characteristics** |  |  |
| **NBV (L) mean ± SD** | 1.46 ± 0.07 | 1.41 ± 0.13 |
| **NWMV (L) mean ± SD** | 0.70 ± 0.04 | 0.73 ± 0.08 |
| **NGMV (L) mean ± SD** | 0.79 ± 0.04 | 0.67 ± 0.09* |
| **Pathological and genetic characteristics** |  |  |
| **Thal phase<sup>47</sup> N** | 9 | 19*** |
| 0/1/2/3/4/5 | 2/3/3/1/0/0 | 0/0/0/1/1/17 |
| **Braak NFT stage<sup>2</sup> N** | 9 | 19*** |
| 0/1/2/3/4/5/6 | 1/7/1/0/0/0/0 | 0/0/0/0/4/8/7 |
| **ABC score<sup>48</sup> N** | 9 | 19 |
| A 0/1/2/3 | 2/6/1/0 | 0/0/0/19*** |
| B 0/1/2/3 | 1/8/0/0 | 0/0/4/15*** |
| C 0/1/2/3 | 9/0/0/0 | 0/0/4/15*** |
| **CAA type<sup>49</sup> N** | 2 | 17 |
| Type 1 / type 2 | 1/1 | 14/3 (82%) |
| **APOE genotype N** | 8 | 19 |
| ε4 non-carrier | 5 (56%) | 7 (37%) |
| ε4 heterozygous | 3 (44%) | 10 (53%) |
| ε4 homozygous | - | 2 (10%) |

Abbreviations: F/M= female/male ratio; SD=standard deviation; N=sample size; L=liter; NBV=normalized brain volume; NWM=normalized white matter volume; NGMV=normalized grey matter volume; NFT=neurofibrillary tangles; CAA=cerebral amyloid angiopathy. \* $p < .05$ , compared to controls. \*\*\* $p < .001$ , compared to controls.

### NbM volume, microstructure and tracts

Both left and right normalized NbM volumes were lower in Alzheimer’s disease donors compared to controls (p=.027 and p=.036 respectively, Fig.3A). Right FA (p=.069) and left MD (p=.478) of the NbM were comparable between groups, but both left NbM FA and right MD were higher in Alzheimer’s disease donors compared to controls (p=.047 and p=.030, respectively, Fig.3B-C). Correlations were found between NbM volume and NbM MD in the right (r= −0.60, p=.001), but not left hemisphere (p=.236) (Fig.3D-E and Supplementary Table 5).

**Figure 3.**
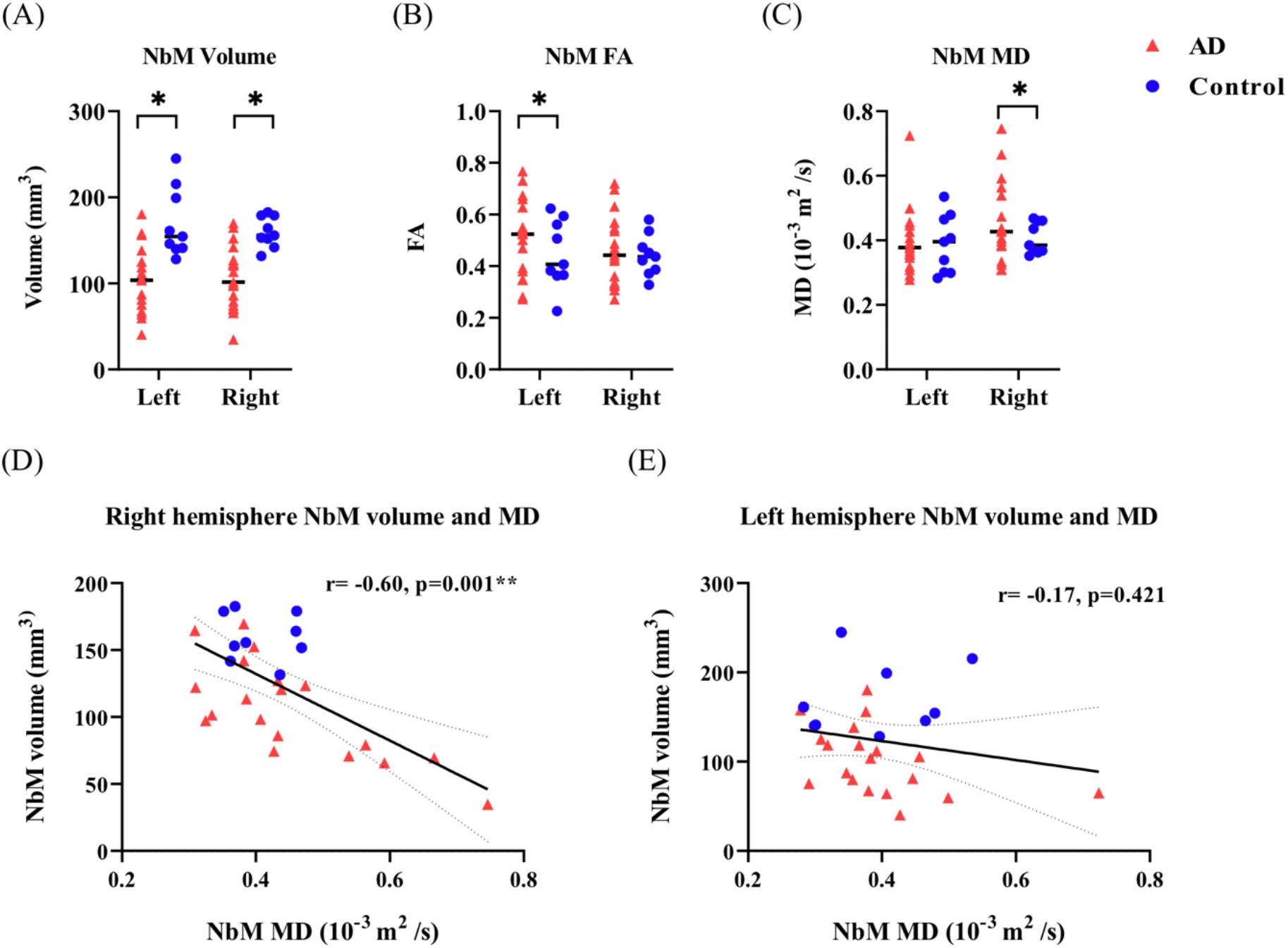
MRI measures of NbM volume, microstructure and their associations. AD cases showed significantly lower NbM volumes in the left (*p*=.027) and right (*p*=.036) hemispheres in (A), higher FA in NbM of the left hemisphere (*p*=.047) in (B), and higher NbM MD in the right hemisphere (*p*=.030) in (C) compared to controls. The NbM volumes significantly correlated with the NbM MD in the right hemisphere (*p*=.001), as shown in (D), however, no significant correlation was found for the left hemisphere (*p*=.421), as shown in (E). For detailed information see Supplementary Table 3. Abbreviations: AD=Alzheimer’s Disease, MD=mean diffusivity, mm=millimeter, s=second, r=rho. \**p*<.05. \*\**p*<.01.

Alzheimer’s disease donors did not show differential NbM tract integrity (FA or MD) to cortical lobes compared to controls (Supplementary Table 4). In the right hemisphere, decreased NbM volume was associated with reduced microstructural integrity of tracts between the NbM and the temporal lobe (FA: r= 0.52, p=.048, FDR-corrected, MD: r=−0.58, p=.018, FDR-corrected), but not to other cortical or cingulate areas (Supplementary Table 5).

### NbM pathological load and ChAT cell density

Alzheimer’s disease donors showed higher pathology load (%area) of Aβ (p=.002) and p-tau (p<.001) than controls (Supplementary Table 6). No significant difference in ChAT cell density was observed between groups (p=.384). This is most likely due to the large variability in both groups, particularly in the AD group (Fig.4). ChAT cell density was not associated with local Aβ (p=.467) or p-tau load (p=.871).

**Figure 4.**
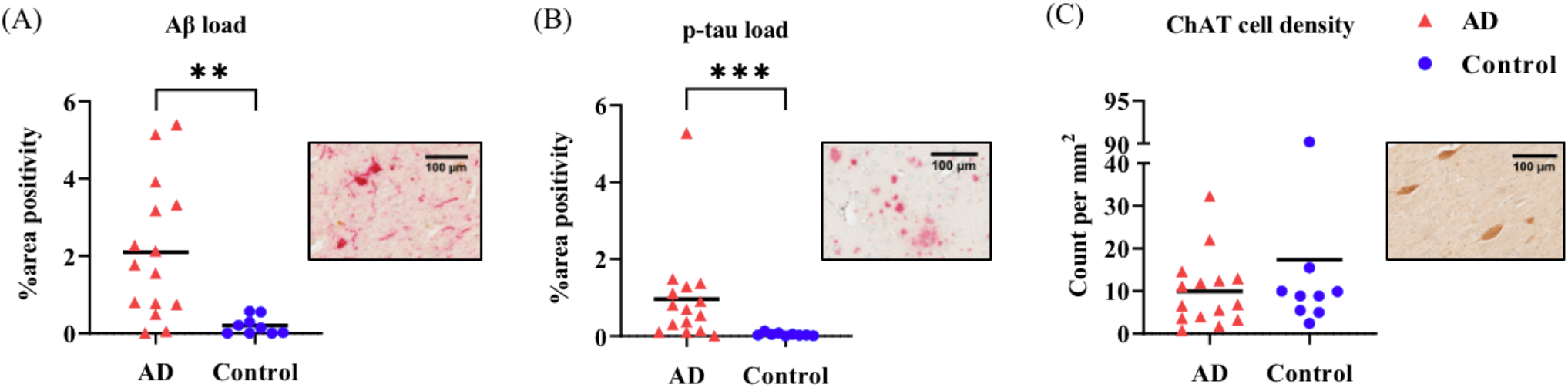
Alzheimer’s disease cases showed higher pathology loads but not ChAT cell density compared to controls. Alzheimer’s disease donors had significantly higher Aβ (*p*=.002) and p-tau (*p*=.0002) load as shown in (A)(B). However, ChAT cell density did not differ to controls, as shown in (C). Of note, some AD donors had very little ChAT cell density, while others had similar counts to controls, illustrating the heterogeneity in our AD cohort. Pathology inserts show the immunoreactivity of Aβ (A), p-tau (B) and ChAT (C). Abbreviations: AD=Alzheimer’s disease, ChAT=choline acetyltransferase, p-tau=phosphorylated tau. Aβ=amyloid-beta. \*\**p*<.01. \*\*\**p*<.001.

### ChAT cell density associated with microstructural integrity of NbM and tracts to the temporal cortex

When assessing MRI and histopathological associations in the whole cohort, MRI-derived NbM MD correlated negatively with ChAT cell density (r= -.49, p=.028), but not with Aβ and p-tau load (p=.670 and p=.249 respectively). NbM volume and FA did not associate with ChAT cell density, Aβ, and p-tau load (all p>.05). (Supplementary Table 7).

When considering the microstructural integrity of tracts between the NbM and cortical lobes, ChAT cell density was negatively associated with MD of tracts to the temporal lobe (β= −0.194, p=.024, FDR-corrected), while no associations were found in other cortical tracts (Fig.5A and Supplementary Table 8). When excluding one control case exhibiting especially high ChAT cell density (90.186 count per mm^2^, falling outside the inner quartile), the association did not survive correction for multiple comparisons (β= −0.182, p=.011, uncorrected; p=.064, FDR-corrected). No significant results were found for NbM tract integrity and Aβ, and p-tau load. Association estimates and p-values of tracts to cingulate and cortical lobes are shown in Supplementary Table 8.

**Figure 5.**
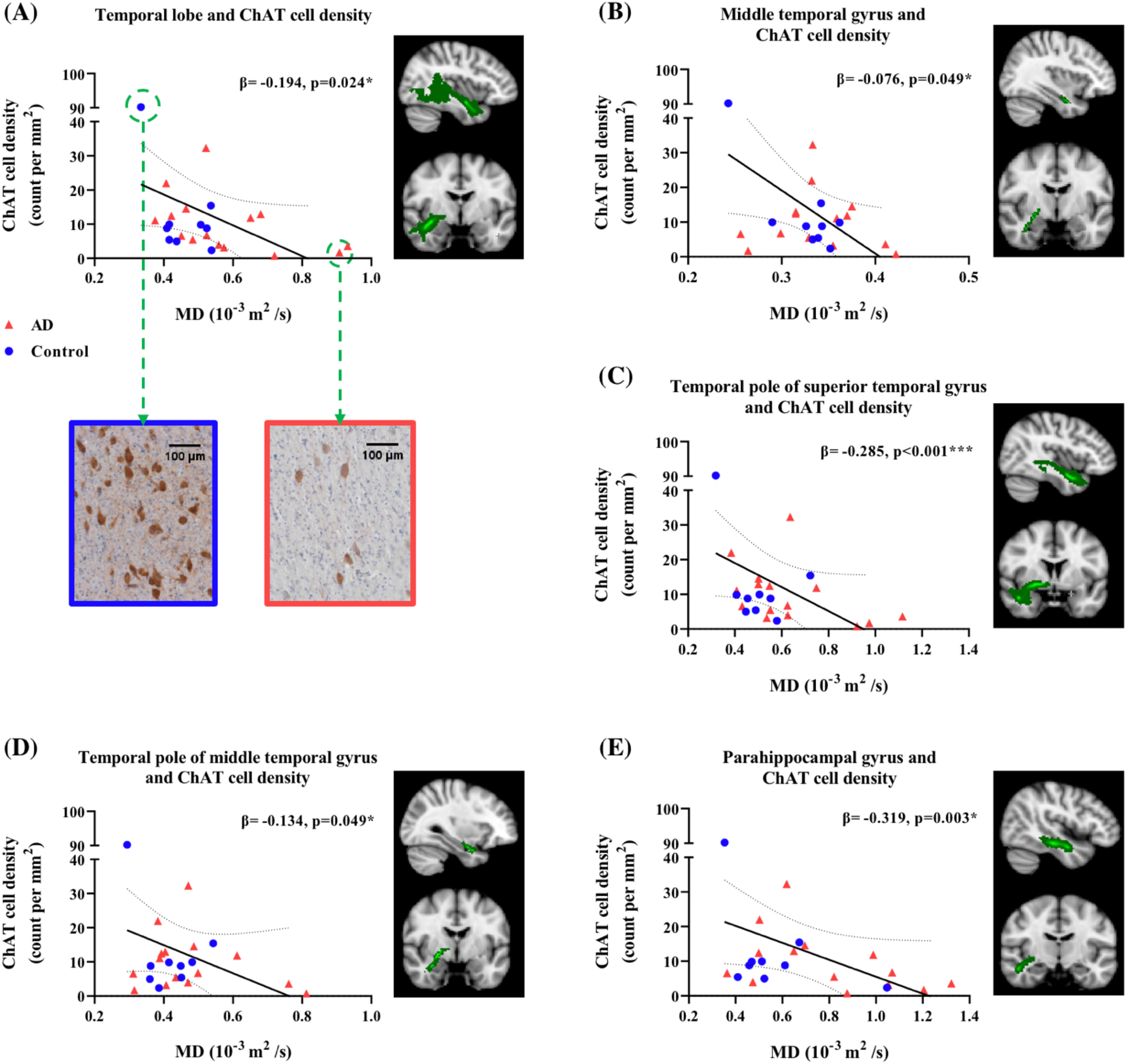
ChAT cell density associated with MD of tracts to the temporal lobe and subregions. ChAT cell density was negatively associated with MD of tracts to temporal lobe, as shown in (A). Pathology inserts show a control case with high ChAT cell density exhibiting low tract MD, and an Alzheimer’s disease case with low ChAT cell density exhibiting high tract MD. Negative associations were also found between ChAT cell density and MD of tracts to temporal subregions: middle temporal gyrus (B), temporal pole of superior and middle temporal gyrus (C)(D), and parahippocampal gyrus. After removal of a case with very high ChAT cell density, the association remained in temporal pole of superior temporal gyrus (β= −0.273, *p*=.003, FDR-corrected) and parahippocampal gyrus (β= −0.316, *p*=.016, FDR-corrected). Abbreviations: AD=Alzheimer’s disease, ChAT=choline acetyltransferase, μm=micron, MD=mean diffusivity, mm=millimeter, s=second. \**p*<.05, FDR-corrected. \*\**p*<.01, FDR-corrected, \*\*\**p*<.001, FDR-corrected.

### ChAT cell density associates with MD in tracts to temporal pole of superior temporal gyrus and parahippocampal gyrus

The above results suggest specific alterations in the tracts to the temporal lobe, of which the decreased tract MD was associated with reduced ChAT cell density. To explore this further, we investigated the tracts to subdivisions of the temporal cortex. As such, ChAT cell density was negatively associated with MD of the tracts to the middle temporal gyrus (β= −0.076, p=.049, FDR-corrected), temporal pole of superior temporal gyrus (β= −0.285, p<.001, FDR corrected), temporal pole of middle temporal gyrus (β= −0.134, p=.049, FDR-corrected), and parahippocampal gyrus (β= −0.319, p=.003, FDR-corrected), as shown in Table 2. When excluding the aforementioned outlier case, the associations between ChAT cell density and tract MD remained significant in tracts to the temporal pole of the superior temporal gyrus and parahippocampal gyrus (respectively β= −0.273, p=.003 and β= −0.316, p=.016, FDR-corrected) (Fig.5).

**Table 2.** Associations between ChAT cell density and the integrity of tracts to regions within the temporal lobe.

|  | Superior<br>temporal<br>gyrus | Middle<br>temporal<br>gyrus | Inferior<br>temporal<br>gyrus | Temporal<br>pole,<br>superior<br>temporal<br>gyrus | Temporal<br>pole, middle<br>temporal<br>gyrus | Parahippocampal<br>gyrus | Hippocampus |
| --- | --- | --- | --- | --- | --- | --- | --- |
| **Tract FA** |  |  |  |  |  |  |  |
| ChAT<br>cell<br>density | p=.883 | p=.391 | p=.872 | p=.124 | p=.872 | $\beta$=0.182<br>p=.027* | p=0.872 |
| **Tract MD** |  |  |  |  |  |  |  |
| ChAT<br>cell<br>density | p=.191 | $\beta$=-0.076<br>p=.049* | $\beta$=-0.073<br>p=.053 | $\beta$=-0.285<br>p<.001*** | $\beta$=-0.134<br>p=.049* | $\beta$=-0.319<br>p=.003** | p=.349 |

Abbreviations: FA=fractional anisotropy, MD=mean diffusivity, ChAT=choline acetyltransferase. \* $p$ <.05, FDR-corrected. \*\* $p$ <.01, FDR-corrected. \*\*\* $p$ <.001, FDR-corrected.

### Exploration in Alzheimer’s disease subtypes

Our Alzheimer’s disease cohort contained amnestic and non-amnestic clinical subtypes; due to this heterogeneity we explored a possible phenotypic effect in our results. Supplementary Table 9 shows the clinical, neuropathological and radiological characteristics of the phenotypes. No significant difference in these characteristics were found between the two subtypes, but both subtypes showed differences compared to controls.

MRI-derived volume of both left and right NbM were lower in the amnestic subtype compared to controls (left: p=.018 and right: p=.029, uncorrected), but no difference between the non-amnestic subtype and controls, as well as between the two subtypes was found. Only tracts between the NbM and temporal lobe showed higher FA in amnestic compared to non-amnestic subtype (p=.036, Bonferroni-corrected).

For histopathological measures, both subtypes had higher p-tau load than controls (amnestic: p=.001 and non-amnestic: p=.018, Bonferroni-corrected), while only the amnestic subtype had higher Aβ load than controls (p=.006, Bonferroni-corrected). No significant difference in pathology loads between subtypes, as well as ChAT cell density between each group was found (Supplementary Table 10).

When examining MRI-histopathology associations in the amnestic subtype, positive associations were found between p-tau load and MRI-derived NbM volume (r=0.75, p=.032), p-tau load and the FA of NbM tracts to the temporal lobe (β= 0.107, p=.018, FDR-corrected), whereas negative associations were found between ChAT cell density and MD of tracts between the NbM and frontal, temporal and parietal lobes (respectively β= −0.295, p=.024; β= −0.306, p=.018; β= −0.355, p=.040, FDR-corrected). In contrast, we did not find any MRI-histopathology associations in non-amnestic subtype (Supplementary Table 11).

## Discussion

In a combined post-mortem *in-situ* MRI and histopathology approach, we investigated the associations between MRI-derived NbM volume, microstructure and trajectories with NbM protein aggregation and cholinergic cell density, in Alzheimer’s disease and non-neurological control donors. We found reduced NbM microstructural integrity to be associated with decreased cholinergic cell density. Furthermore, this decrease in the cholinergic cell density in the NbM was associated with altered integrity in white matter tracts between the NbM and regions within the temporal cortex. Interestingly, this was specifically the case for amnestic rather than non-amnestic Alzheimer’s disease donors.

In our study, post-mortem NbM MRI volume loss was found in Alzheimer’s disease compared to control donors, which has also been described previously in the *in vivo* literature^11,50^. We did not find a group difference in the integrity of tracts between the NbM and the frontal and temporal lobe. Although not previously addressed in *in vivo* neuroimaging studies, histological studies have shown disrupted cholinergic pathways and cholinergic denervation to the frontal and temporal lobe in Alzheimer’s disease^51,52^. The reason why we did not find such difference may be due to the heterogeneity of our Alzheimer’s disease cohort, as we did find differences in tract integrity between the NbM and temporal cortex between amnestic and non-amnestic subtypes. We also showed that NbM atrophy was associated with local decreased microstructural integrity, similar to the association found in an *in vivo* MRI study in Alzheimer’s disease^16^. Moreover, we found increased NbM MD to be associated with lower cholinergic cell density, suggesting that MRI-derived MD may be a sensitive marker for cholinergic degeneration^53–57^.

We did not find an absolute reduction in cholinergic cell density in Alzheimer’s disease compared to control donors, a finding that is both consistent and contradictory to the literature^5,8,58–60^. It is suggested that these inconsistent results reflect the large heterogeneity in cholinergic cell loss within subsectors of the NbM^4^. There appears to be a caudorostral gradient of NbM neuronal loss in Alzheimer’s disease, with the posterior sector being the most severely affected^4,8,61,62^. We identified our tissue sections as part of the pre-anterior/ anteroromediate/ antero-intermediate NbM^4,41,43^, which could explain why we did not find a group difference in cholinergic cell density.

Both NbM Aβ and p-tau load were higher in Alzheimer’s disease donors of our cohort. However, we found no associations between local Aβ, p-tau load and MRI outcome measures. Indeed previous studies only showed an association between cortical (and not NbM) Aβ and MRI-derived NbM volume^63^. Results for tau are more incongruent, one study did not find an association between MRI-derived NbM volume and Braak NFT stage^64^, while another study found an association between NbM volume and p-tau levels in cerebrospinal fluid^65^. A possible explanation could be found in our exploration in Alzheimer’s disease phenotypes, in which only amnestic Alzheimer’s disease donors showed an association between NbM MRI volume and p-tau. As this association was made with a very small sample size, this should be futher explored in a larger cohort with a variety of phenotypes^1,41^.

We found decreased ChAT cell density associated with an increased MD of tracts to the temporal lobe, specifically to the temporal pole of superior temporal gyrus and parahippocampal gyrus, suggesting that effects of cholinergic degeneration on cholinergic pathways can be captured with diffusion MRI. The lateral cholinergic pathways of the NbM innervates the superior, middle and inferior temporal gyrus and parahippocampal gyrus, and is responsible for cortical cholinergic signaling in the temporal lobe^19,66^. Cholinergic denervation may leads to reduced acetylcholine in the medial temporal cortex, and decreases the spiking activity in cholinergic neurons, which in turn decreasing activation of cholinergic receptors, undermining memory formation in Alzheimer’s disease^9,21,67–69^. Specifically parahippocampal acetylcholine activity plays a role in mediating memory encoding and consolidation^67,70^. This shows the crucial role of temporal cholinergic projections in supporting memory-related circuitry in the medial temporal lobe, which is clinically relevant in amnestic Alzheimer’s disease^12,13,71^.

The cholinergic system is differentially affected in clinically-defined Alzheimer’s disease subtypes^72^. Memory deficits in amnestic Alzheimer’s disease have been linked to cholinergic dysfunction^5^, whereas non-amnestic Alzheimer’s disease with atypical symptoms, such as visual hallucination and aphasia, has been linked with focal pathological change in specific brain areas^73,74^. We explored the cholinergic integrity among the subtypes in our cohort and found cholinergic alterations only within the amnestic subtype. Compared to controls, MRI-derived volume reduction and higher pathological burden was found in the NbM of amnestic donors, while no difference was shown in the non-amnestic donors. Furthermore, decreased ChAT cell density was associated with increased MD of tracts between the NbM and frontal, temporal and parietal lobes only within amnestic Alzheimer’s disease. Altogether, cholinergic transmission to the neocortex may be predominantly interrupted in the amnestic subtype, as illustrated by altered integrity in NbM tracts and memory-related symptoms. Moreover, p-tau accumulation in the NbM may play a role in this cholinergic disruption, especially to the temporal lobe^75^.

More bridging evidence between neuroimaging and histopathology is needed to support our results and further validate and identify useful imaging (bio)markers and potential posttreatment auxiliary in Alzheimer’s disease^76–78^. A few limitations of our study need to be addressed. First our small sample size, a further validation in a larger cohort is required to better account for the clinical and pathological heterogeneity and to be able to differentiate cholinergic projections to cortical subregions in Alzheimer’s disease phenotypes. In addition, our NbM tissue blocks were retrospectively collected from the brain bank and not directly at the autopsy where the dissecting plain for the NbM block can be derived more consistently across cases to reduce the variability in NbM subsectors. Furthermore, recent studies have also raised attention to co-morbid pathology in Alzheimer’s disease, such as TAR DNA-binding protein 43 and Lewy body pathology^79,80^. Future research is encouraged to account for alternative secondary pathology to better understand the pathological mechanisms underlying cholinergic degeneration in Alzheimer’s disease and related disorders.

## Conclusion

The present study investigated cholinergic degeneration within the NbM and its structural projections to cortical regions with combined post-mortem MRI and histopathology. Alzheimer’s disease donors showed altered reduced NbM volume and altered NbM integrity. In addition, the decline in cholinergic cell density was associated with microstructural alterations in the NbM and its projections, specifically to the temporal cortex. Together, these findings indicate that the pathological alteration is not limited within the NbM but can also be captured in its cortical projections, especially for patients with an amnestic Alzheimer’s disease phenotype. The current study reveals the association between MRI and histopathology of the cholinergic system in Alzheimer’s disease, which provides valuable insights in the pathophysiology underlying *in-vivo* imaging (bio)markers.

## Abbreviations

NbM: nucleus basalis of Meynert
FA: fractional anisotropy
MD: mean diffusivity
p-tau: phosphorylated-tau
Aβ: amyloid-beta
ChAT: choline acetyltransferase

## Acknowledgements

We would like to thank all brain donors and their next of kin for brain donation, the Netherlands Brain Bank, the Normal Aging Brain Collection Amsterdam (NABCA) autopsy team.

