## Supplementary materials for "Structural (dys)connectivity associates with cholinergic cell density of the nucleus basalis of Meynert in Alzheimer’s disease"

**Methods**

*MRI protocol*

T1-weighted images (T1w) were acquired using a sagittal 3D T1-weighted fast spoiled gradient echo sequence with the following parameters: repetition time (TR)/echo time (TE)/ inversion time (TI) = 7/3/450 ms, flip angle = 15°, slice thickness= 1 mm, in-plane resolution= 1.0 x 1.0 $mm^2$. A sagittal 3D fluid attenuation inversion recovery was acquired with TR/TE/TI= 8000/130/2000-2250 ms, slice thickness 1.2 mm, in-plane resolution= 1.11 × 1.11 $mm^2$. TI was optimized per case to account for post-mortem delay and temperature differences leading to variable cerebrospinal fluid suppression. Diffusion-weighted imaging was acquired by axial 2D echo-planar imaging with diffusion gradients applied in 30 non-collinear directions, TR/TE= 7400/92 ms, slice thickness 2.0 mm, in-plane resolution=2.0 x 2.0 $mm^2$, and b=1000 $s/mm^2$. To allow for offline distortion correction of the images, 5 b0 images were acquired using the same sequence parameters.

*Immunohistochemistry for cholinergic neurons and Aβ and p-tau in NbM sections*

Paraffin-embedded tissue blocks of the NbM were cut at 20 µm (Leica Microtome) for 30 serial sections and mounted onto glass slides (Thermo Scientific, USA) for subsequent immunohistochemistry with Choline Acetyltransferase (ChAT), amyloid-beta (Aβ) and phosphorylated tau (p-tau) pathology (see Supplementary Table 2 for detailed information on antibodies).

From the 30 sections that were sliced, starting from the first section with visible anterior commissure and substantia innominata, three slides were taken at a distance of 200 um (e.g., every 1st, 11th and 21th, every 2nd, 12th and 22th, and every 3rd, 13th and 23rd slide) to account for variability within the NbM. These selected sections were deparaffinized and rehydrated in a graded series of xylene and ethanol. Subsequently, the sections underwent antigen retrieval in citrate buffer (pH 6.0) at a temperature of 95°C. The sections were first blocked for endogenous peroxidase by immersing the sections in 1% of $H_2O_2$ , in tris-buffered saline (TBS, pH 7.4) for thirty minutes. Subsequently the sections were blocked with 0.1% Triton and 3% normal donkey serum. The double staining of ChAT and Aβ sections underwent additional second antigen retrieval using 80% formic acid for five minutes in the fume hood before the first block. All sections were incubated with primary antibodies, ChAT, ChAT/p-tau (AT8) and ChAT/Aβ (6F/3D), diluted in TBS for two nights at 4°C, followed by Immpress (Vector, California, United State) detection. Finally, ChAT was visualized with 3,3'-Diaminobenzidine (DAB, Dako, Glostrup, Denmark) imidazole (50 mg DAB, 350 mg Imidazole and 30 uL of $H_2O_2$ per 100 mL of Tris-HCl 30mM, pH 7.6), AT8 and 6F/3D were visualized using liquid permanent red followed by counter-staining with haematoxylin, and mounting with Entellan.

**Supplementary Table 1. List of automated anatomical labeling (AAL) regions in each lobe.**

| **Frontal lobe** | **Temporal lobe** | **Parietal lobe** | **Occipital lobe** | **Cingulum** | **Insula** |
|---|---|---|---|---|---|
| Inferior frontal gyrus, pars orbitalis | Heschl's gyrus | Postcentral gyrus | Superior occipital gyrus | Anterior cingulate & paracingulate gyri | |
| Superior frontal gyrus, dorsolateral | Superior temporal gyrus | Superior parietal gyrus | Middle occipital gyrus | Middle cingulate & paracingulate gyri | |
| Middle frontal gyrus | Middle temporal gyrus | Inferior parietal gyrus, excluding supramarginal and angular gyri | Inferior occipital gyrus | Posterior cingulate gyrus | |
| Inferior frontal gyrus, opercular part | Inferior temporal gyrus | SupraMarginal gyrus | Calcarine fissure and surrounding cortex | | |
| Inferior frontal gyrus, triangular part | Temporal pole: superior temporal gyrus | Angular gyrus | Cuneus | | |
| Superior frontal gyrus, medial | Temporal pole: middle temporal gyrus | Precuneus | Lingual gyrus | | |
| Supplementary motor area | ParaHippocampal gyrus | | | | |
| Precentral gyrus | Hippocampus | | | | |

Note: The AAL regions in each lobe applied to both left and right hemisphere[1]. The orbital part of superior and middle frontal lobe, rectus, olfactory cortex, paracentral lobule and rolandic operum in the frontal lobe, along with fusiform in the temporal lobe were excluded due to their susceptibility to DWI distortions.

**Supplementary Table 2. Information on primary antibodies.**

| Antibody | Antigen | Species | Origin details | Dilution | Incubation time | Antigen retrieval | Detection method |
|---|---|---|---|---|---|---|---|
| **ChAT** | ChAT | Goat IgG | Chemicon, Darmstadt, Germany | 1:200 | 4°C o.n. | Autoclave (pH 6.0, 10 min) | Impress |
| **p-tau, clone AT8** | Tau phosphorylated at Ser202 and Thr205 | Mouse IgG1 | ThermoFisher, Pittsburgh, USA | 1:2000 | 4°C o.n. | Autoclave (pH 6.0, 10 min) | Impress |
| **Aβ, clone 6F/3D** | Aβ amino acid sequence 8-17 and additional C-terminal cysteine coupled to keyhole limpet hemocyanin | Mouse IgG2b | Dako, Glostrup, Denmark | 1:2000 | 4°C o.n. | Autoclave (pH 6.0, 10 min) | Impress |

Abbreviations: ChAT=choline acetyltransferase, p-tau=phosphorylated tau, Aβ=amyloid-beta, o.n.=overnight, pH=potential of hydrogen.

### Supplementary Table 3. Detailed donor's information

| Clinical characteristics | | | | | | | | Radiological characteristics | | | | | Pathological, genetic and cognitive characteristics | | | |
|---|---|---|---|---|---|---|---|---|---|---|---|---|---|---|---|---|
| Case # | Group | AD Phenotype | Gender | Disease duration | Age at death | Cause of death | PMD (hr:min) | NBV (L) | NWMV (L) | NGMW (L) | NbM_R (mL) | NbM_L (mL) | Thal phase[2] | Braak NFT stage[3] | ABC[4] | *APOE* |
| 1 | AD | Amnestic | M | 2 | 60 | Euthanasia | 8:35 | 1.551 | 0.787 | 0.763 | 152.36 | 156.05 | 5 | 6 | A3 B3 C3 | 33 |
| 2 | AD | Amnestic | M | 6 | 68 | Euthanasia | 9:15 | 1.429 | 0.711 | 0.718 | 123.39 | 118.59 | 5 | 5 | A3 B3 C3 | 33 |
| 3 | AD | Amnestic | M | 11 | 69 | Respiratory failure due to pulmonary infection | 11:55 | 1.664 | 0.971 | 0.693 | 86.09 | 87.30 | 5 | 5 | A3 B3 C3 | 34 |
| 4 | AD | Non-amnestic | M | 7 | 65 | Cardiac arrest | 7:50 | 1.505 | 0.732 | 0.774 | 113.37 | 125.23 | 4 | 5 | A3 B3 C3 | 33 |
| 5 | AD | Non-amnestic | M | NA | 59 | Euthanasia | 6:30 | 1.581 | 0.777 | 0.804 | 74.34 | 67.33 | 5 | 5 | A3 B3 C3 | 44 |
| 6 | AD | Non-amnestic | F | 4 | 78 | Cachexia and dehydration | 7:30 | 1.359 | 0.698 | 0.661 | 127.08 | 111.95 | 5 | 5 | A3 B3 C3 | 34 |
| 7 | AD | Typical | M | 13 | 84 | Euthanasia | 5:53 | 1.434 | 0.711 | 0.723 | 98.28 | 138.37 | 5 | 4 | A3 B2 C2 | 34 |
| 8 | AD | Non-amnestic | M | 8 | 62 | Palliative sedation | 8:15 | 1.155 | 0.623 | 0.532 | 122.14 | 118.40 | 5 | 6 | A3 B3 C3 | 34 |
| 9 | AD | Non-amnestic | M | 5 | 37 | Euthanasia | 11:11 | 1.510 | 0.779 | 0.732 | 142.03 | 103.79 | 5 | 6 | A3 B3 C3 | 23 |
| 10 | AD | Amnestic | F | 7 | 80 | Euthanasia | 7:05 | 1.435 | 0.725 | 0.709 | 164.65 | 157.99 | 5 | 4 | A3 B2 C2 | 33 |
| 11 | AD | Non-amnestic | M | 9 | 67 | Cachexia and dehydration | 6:35 | 1.254 | 0.671 | 0.583 | 120.60 | 105.38 | 5 | 6 | A3 B3 C3 | 34 |
| 12 | AD | Non-amnestic | M | 4 | 77 | Euthanasia | 7:00 | 1.542 | 0.769 | 0.774 | 169.63 | 180.32 | 5 | 4 | A3 B2 C2 | 34 |
| 13 | AD | Amnestic | M | 5 | 53 | Palliative sedation | 9:00 | 1.264 | 0.668 | 0.596 | 79.11 | 81.11 | 5 | 6 | A3 B3 C3 | 33 |
| 14 | AD | Amnestic | M | 12 | 64 | Cachexia and dehydration | 7:55 | 1.251 | 0.683 | 0.568 | 65.70 | 59.72 | 5 | 6 | A3 B3 C3 | 34 |
| 15 | AD | Non-amnestic | M | 3 | 59 | Dysphagia | 5:35 | 1.321 | 0.676 | 0.645 | 69.42 | 64.08 | 5 | 5 | A3 B3 C3 | 34 |
| 16 | AD | Non-amnestic | M | 10 | 73 | Cachexia and dehydration | 7:20 | 1.366 | 0.873 | 0.493 | 34.67 | 40.44 | 5 | 5 | A3 B3 C3 | 34 |
| 17 | AD | Amnestic | M | 23 | 84 | Euthanasia | 8:35 | 1.254 | 0.642 | 0.612 | 70.99 | 64.97 | 3 | 4 | A2 B2 C2 | 33 |

| | | | | | | | | | | | | | | | | |
|---|---|---|---|---|---|---|---|---|---|---|---|---|---|---|---|---|
| 18 | AD | Amnestic | M | 10 | 77 | Suicide by drugs | 9:05 | 1.400 | 0.728 | 0.672 | 101.43 | 75.39 | 5 | 6 | A3 B3 C3 | 44 |
| 19 | AD | Amnestic | M | 7 | 65 | Euthanasia | 9:20 | 1.464 | 0.697 | 0.767 | 97.18 | 79.96 | 5 | 5 | A3 B3 C3 | 34 |
| 20 | Control | NA | M | NA | 68 | Euthanasia | 8:30 | 1.609 | 0.758 | 0.851 | 153.09 | 161.08 | 2 | 1 | A1 B1 C0 | 34 |
| 21 | Control | NA | F | NA | 69 | Pulmonary embolism | 12:40 | 1.375 | 0.671 | 0.704 | 182.56 | 244.86 | 1 | 1 | A1 B1 C0 | 33 |
| 22 | Control | NA | M | NA | 59 | Euthanasia | 8:00 | 1.494 | 0.730 | 0.765 | 151.78 | 154.37 | 2 | 1 | A1 B1 C0 | 34 |
| 23 | Control | NA | F | NA | 78 | Unknown | 5:25 | 1.516 | 0.738 | 0.777 | 164.08 | 199.24 | 1 | 1 | A1 B1 C0 | 33 |
| 24 | Control | NA | F | NA | 59 | Euthanasia | 8:10 | 1.460 | 0.677 | 0.783 | 179.02 | 215.40 | 0 | 0 | A0 B0 C0 | 33 |
| 25 | Control | NA | F | NA | 71 | Lung carcinoma | 6:40 | 1.477 | 0.708 | 0.769 | 178.95 | 141.12 | 2 | 1 | A1 B1 C0 | 34 |
| 26 | Control | NA | M | NA | 72 | Hypovolemic shock | 8:23 | 1.424 | 0.683 | 0.742 | 155.62 | 140.06 | 1 | 1 | A1 B1 C0 | Unknown |
| 27 | Control | NA | F | NA | 87 | Urinary tract infection | 8:35 | 1.367 | 0.652 | 0.715 | 131.62 | 145.92 | 0 | 1 | A0 B1 C0 | Unknown |
| 28 | Control | NA | M | NA | 74 | Metastatic neuroendocrine tumor colon | 16:00 | 1.516 | 0.715 | 0.801 | 141.83 | 128.06 | 2 | 0 | A1 B0 C0 | Unknown |

Abbreviations: AD= Alzheimer's disease, PMD=post-mortem delay, NBV=normalized brain volume, NWMV=normalized white matter volume, NGMV=normalized grey matter volume, L=liter, F/M=female/male, NFT=neurofibrillary tangle, NA=not applicable.

**Supplementary Table 4. Mean and standard deviation of tract FA and MD to cortical lobes in AD and controls.**

| | AD (n=19) | Controls (n=9) | p-values |
|---|---|---|---|
| | **Right hemisphere tract FA** | | |
| **Cingulum** | 0.464±0.100 | 0.460±0.080 | ns |
| **Frontal lobe** | 0.355±0.056 | 0.459±0.097 | ns |
| **Temporal lobe** | 0.371±0.059 | 0.480±0.070 | ns |
| **Parietal lobe** | 0.464±0.036 | 0.358±0.078 | ns |
| **Occipital lobe** | 0.483±0.051 | 0.342±0.052 | ns |
| **Insula** | 0.326±0.073 | 0.525±0.103 | ns |
| | **Right hemisphere tract MD** | | |
| **Cingulum** | 0.457±0.224 | 0.471±0.126 | ns |
| **Frontal lobe** | 0.640±0.164 | 0.459±0.097 | ns |
| **Temporal lobe** | 0.583±0.156 | 0.480±0.070 | ns |
| **Parietal lobe** | 0.425±0.151 | 0.358±0.078 | ns |
| **Occipital lobe** | 0.400±0.103 | 0.342±0.052 | ns |
| **Insula** | 0.709±0.229 | 0.525±0.103 | ns |

| | AD (n=18) | Controls (n=9) | p-values |
|---|---|---|---|
| | **Left hemisphere tract FA** | | |
| **Cingulum** | 0.517±0.097 | 0.470±0.053 | ns |
| **Frontal lobe** | 0.375±0.086 | 0.381±0.082 | ns |
| **Temporal lobe** | 0.370±0.067 | 0.381±0.046 | ns |
| **Parietal lobe** | 0.498±0.045 | 0.515±0.019 | ns |
| **Occipital lobe** | 0.516±0.047 | 0.505±0.029 | ns |
| **Insula** | 0.313±0.076 | 0.304±0.050 | ns |
| | **Left hemisphere tract MD** | | |
| **Cingulum** | 0.385±0.079 | 0.396±0.126 | ns |
| **Frontal lobe** | 0.666±0.337 | 0.456±0.139 | ns |
| **Temporal lobe** | 0.680±0.291 | 0.524±0.083 | ns |
| **Parietal lobe** | 0.360±0.096 | 0.337±0.062 | ns |
| **Occipital lobe** | 0.359±0.071 | 0.334±0.070 | ns |
| **Insula** | 0.746±0.289 | 0.563±0.158 | ns |

Group comparisons between Alzheimer's disease donors and controls were conducted using GLM and corrected for multiple comparisons. MD reported in MD ($10^{-3}$ $m^2/s$). Abbreviations: AD=Alzheimer's disease, FA=fractional anisotropy, MD=mean diffusivity, ns=not significant.

**Supplementary Table 5. Correlations between NbM volumes and the integrity of NbM and cortical tracts.**

| | Left hemisphere NbM volume | Right hemisphere NbM volume |
|---|---|---|
| **NbM FA** | r=0.22, p= 0.299 | r=0.25, p= 0.236 |
| **Cingulum tract FA** | r=0.10, p= 0.884 | r=0.19, p=0.526 |
| **Frontal tract FA** | r= 0.05, p= 0.884 | r= 0.24, p= 0.402 |
| **Temporal tract FA** | r= 0.03, p= 0.884 | r= 0.52, p= 0.048* |
| **Parietal tract FA** | r= 0.20, p= 0.884 | r= 0.25, p= 0.402 |
| **Occipital tract FA** | r= -0.17, p= 0.884 | r= 0.08, p= 0.692 |
| **Insula tract FA** | r= 0.38, p= 0.564 | r= 0.27, p= 0.402 |
| **NbM MD** | r= -0.17, p= 0.421 | r= -0.60, p= 0.001 |
| **Cingulum tract MD** | r= -0.17, p=0.643 | r= -0.39, p= 0.102 |
| **Frontal tract MD** | r= -0.23, p=0.643 | r= -0.46, p= 0.023^ |
| **Temporal tract MD** | r= 0.01, p= 0.947 | r= -0.58, p=0.018* |
| **Parietal tract MD** | r= 0.18, p=0.643 | r= -0.43, p= 0.086 |
| **Occipital tract MD** | r= 0.16, p=0.643 | r= -0.40, p= 0.086 |
| **Insula tract MD** | r= -0.30, p=0.643 | r= -0.35, p= 0.102 |

Correlations were conducted using partial correlation with age and PMD as covariates. Abbreviations: FA=fractional anisotropy, MD= mean diffusivity, NbM=nucleus basalis of Meynert, r=rho. *p<.05, FDR-corrected. ^p<.05, uncorrected.

**Supplementary Table 6. Means and standard deviations of histopathological measures in AD and controls.**

| | AD (n=15) | Controls (n=9) | p-values |
|---|---|---|---|
| **ChAT cell density** | 9.92±8.45 | 17.31±27.58 | p=0.434 |
| **p-tau load (%area)** | 0.96±1.29 | 0.04±0.04 | p=0.0002*** |
| **Aβ load (%area)** | 2.10±1.75 | 0.20±0.23 | p=0.002** |

Group comparisons between AD and controls were conducted using GLM. Abbreviations: AD=Alzheimer's disease, ChAT=choline acetyltransferase, p-tau=phosphorylated-tau, Aβ=amyloid-beta. **p<.01. ***p<.001.

**Supplementary Table 7. Correlations between MRI-derived NbM volume and integrity with histopathological measures.**

| | NbM volume | NbM FA | NbM MD |
|---|---|---|---|
| **ChAT cell density** | r=0.37<br>p=0.112 | r=0.27<br>p=0.246 | r=-0.49<br>p=0.028* |
| **Aβ load (%area)** | r=-0.20<br>P=0.374 | r=0.09<br>p=0.715 | r=0.04<br>p=0.882 |
| **p-tau load (%area)** | r=-0.18<br>p=0.437 | r=0.17<br>p=0.464 | r=-0.12<br>p=0.613 |

Pearson correlation was used to assess the correlations with age, PMD and gender as covariates. Abbreviations: NbM=nucleus basalis of Meynert, r=rho, FA=fractional anisotropy, MD= mean diffusivity, ChAT=cholinergic acetyltransferase, p-tau=phosphorylated-tau, Aβ=amyloid-beta. *p<.05.

**Supplementary Table 8. Associations between NbM tract integrity and histopathological measures.**

| | Tract FA | | | | | |
|---|---|---|---|---|---|---|
| | **Cingulum** | **Frontal lobe** | **Temporal lobe** | **Parietal lobe** | **Occipital lobe** | **Insula** |
| **ChAT cell density** | p=0.896 | p=0.896 | p=0.896 | p=0.896 | p=0.896 | p=0.896 |
| **Aβ load (%area)** | p=0.878 | p=0.878 | p=0.878 | p=0.878 | p=0.878 | p=0.878 |
| **p-tau load (%area)** | p=0.863 | p=0.824 | p=0.824 | p=0.863 | p=0.863 | p=0.824 |
| | **Tract MD** | | | | | |
| | **Cingulum** | **Frontal lobe** | **Temporal lobe** | **Parietal lobe** | **Occipital lobe** | **Insula** |
| **ChAT cell density** | p=0.272 | p=0.189 | **β= -0.194, p=0.024*** | β= -0.246, p=0.057 | p=0.142 | p=0.369 |
| **Aβ load (%area)** | p=0.501 | p=0.811 | p=0.811 | p=0.722 | p=0.162 | p=0.811 |
| **p-tau load (%area)** | p=0.958 | p=0.870 | p=0.920 | p=0.870 | p=0.870 | p=0.920 |

Abbreviations: FA=fractional anisotropy, MD=mean diffusivity, ChAT=choline acetyltransferase. Aβ= Amyloid-β, p-tau=phosphorylated-tau
*p<.05, FDR-corrected.

**Supplementary Table 9. Donor characteristics of amnestic and non-amnestic AD.**

| | Controls | Amnestic AD | Non-amnestic AD |
|---|---|---|---|
| | | **Clinical characteristics** | |
| **N (phenotype)** | 9 | 10 | 9 (6 B/D, 3 PCA) |
| **Gender F/M (%M)** | 4/5 (56%) | 2/8 (80%)* | 2/7 (20%)* |
| **Age at death**<br>years, mean ± SD | 70.8 ± 8.8 | 70.4 ± 10.5 | 65.4 ± 12.4 |
| **Disease duration**<br>years, mean ± SD | | 7.5 ± 3.8 | 6.9 ± 3.1 |
| **PMD,** mean (hr:min) ± SD (hr) | 9:10 ± 3 | 8:19 ± 2 | 7:13 ± 2 |
| | | **Radiological characteristics** | |
| **NBV (L**) mean ± SD | 1.46 ± 0.07 | 1.42 ± 0.13 | 1.40 ± 0.14 |
| **NGMV (L)** mean ± SD | 0.70 ± 0.04 | 0.68 ± 0.07** | 0.67 ± 0.11* |
| **NWMV (L)** mean ± SD | 0.79 ± 0.04 | 0.73 ± 0.09 | 0.73 ± 0.08 |
| | | **Pathological and genetic characteristics** | |
| **Thal phase N**<br>0/1/2/3/4/5 | 9<br>2/3/3/1/0/0 | 10***<br>0/0/0/1/0/9 | 9***<br>0/0/0/0/1/8 |
| **Braak NFT stage N**<br>0/1/2/3/4/5/6 | 9<br>1/7/1/0/0/0/0 | 10***<br>0/0/0/0/3/3/4 | 9***<br>0/0/0/0/1/5/3 |
| **ABC score N** | 9 | 10** | 9** |

| | | | |
|---|---|---|---|
| A 0/1/2/3 | 2/6/1/0 | 0/0/0/10 | 0/0/0/9 |
| B 0/1/2/3 | 1/8/0/0 | 0/0/3/7 | 0/0/1/8 |
| C 0/1/2/3 | 9/0/0/0 | 0/0/3/7 | 0/0/1/8 |
| **CAA type N** | 2 | 9 | 9 |
| Type 1 / type 2 | 1/1 | 7/2 (78%) | 8/1 (89%) |
| **APOE genotype N** | 8 | 10 | 9 |
| ε4 non-carrier | 5 (56%) | 5 (50%) | 2 (22%) |
| ε4 heterozygous | 3 (44%) | 4 (40%) | 6 (67%) |
| ε4 homozygous | - | 1 (10%) | 1 (11%) |

Abbreviations: AD=Alzheimer's disease, B/D=behavioral/dysexecutive variant; PCA=posterior cortical atrophy; F/M= female/male ratio; SD=standard deviation; N=sample size; L=liter; NBV=normalized brain volume; NWM=normalized white matter volume; NGMV=normalized grey matter volume; NFT=neurofibrillary tangles; CAA=cerebral amyloid angiopathy. *p<.05, compared to controls. **p<.01, compared to controls. ***p=.001, compared to controls.

**Supplementary Table 10. Group comparisons of MRI and histopathological measures between amnestic and non-amnestic AD phenotypes.**

| | Controls (n=9) | Amnestic AD (n=10) | Non-amnestic AD (n=9) | Each subtype compares to controls and p-values | Compare between subtypes and p-values |
|---|---|---|---|---|---|
| | | | **NbM volumes** | | |
| **Right** | 159.8±17.7 | 103.9±33.3 | 108.4±41.4 | Controls>Amnestic, p=.018^ | ns |
| **Left** | 170.0±40.2 | 101.9±37.5 | 109.6±36.3 | Controls>Amnestic, p=.029^ | ns |
| | | | **NbM FA** | | |
| **Right** | 0.443±0.079 | 0.437±0.102 | 0.400±0.106 | ns | ns |
| **Left** | 0.448±0.131 | 0.399±0.132 | 0.444±0.097 | ns | ns |
| | | | **NbM MD** | | |
| **Right** | 0.407±0.048 | 0.437±0.102 | 0.463±0.144 | ns | ns |
| **Left** | 0.389±0.090 | 0.399±0.132 | 0.384±0.041 | ns | ns |
| | | | **Right hemisphere tract FA** | | |
| **Cingulum** | 0.460±0.080 | 0.488±0.049 | 0.440±0.133 | ns | ns |
| **Frontal lobe** | 0.459±0.097 | 0.355±0.054 | 0.356±0.062 | ns | ns |
| **Temporal lobe** | 0.480±0.070 | 0.397±0.052 | 0.343±0.056 | Amnestic>Controls, p=.063^ | Amnestic>Non-amnestic, p=.036* |
| **Parietal lobe** | 0.358±0.078 | 0.452±0.033 | 0.482±0.035 | ns | ns |
| **Occipital lobe** | 0.342±0.052 | 0.483±0.029 | 0.483±0.075 | ns | ns |
| **Insula** | 0.525±0.103 | 0.360±0.077 | 0.288±0.049 | ns | Amnestic>Non-amnestic, p=.026^ |
| | | | **Right hemisphere tract MD** | | |
| **Cingulum** | 0.471±0.126 | 0.491±0.294 | 0.423±0.135 | ns | ns |
| **Frontal lobe** | 0.459±0.097 | 0.666±0.149 | 0.613±0.183 | ns | ns |
| **Temporal lobe** | 0.480±0.070 | 0.537±0.155 | 0.634±0.150 | ns | ns |
| **Parietal lobe** | 0.358±0.078 | 0.463±0.182 | 0.372±0.075 | ns | ns |
| **Occipital lobe** | 0.342±0.052 | 0.422±0.109 | 0.369±0.091 | ns | ns |
| **Insula** | 0.525±0.103 | 0.621±0.184 | 0.807±0.243 | ns | Amnestic>Non-amnestic, p=.038^ |

| | Controls (n=9) | Amnestic AD (n=8) | Non-amnestic AD (n=7) | | |
|---|---|---|---|---|---|
| | | | **Histopathological load** | | |
| **ChAT cell density** | 17.31±27.58 | 9.73±6.62 | 10.14±10.75 | ns | ns |
| **p-tau load (%area)** | 0.04±0.04 | 1.26±1.69 | 0.63±0.58 | Amnestic>Controls, p=.001**; Non-amnestic>Controls, p=.018* | ns |
| **Aβ load (%area)** | 0.20±0.23 | 2.60±1.92 | 1.53±1.46 | Amnestic>Controls, p=.006** | ns |

Group comparisons were performed with general linear model with age and post-mortem delay as covariates. MD reported in MD ($10^{-3}$ $m^2/s$). Abbreviations: AD=Alzheimer's disease, NbM=nucleus basalis of Meynert, FA=fractional anisotropy, MD=mean diffusivity, ChAT=cholinergic acetyltransferase, p-tau=phosphorylated-tau, Aβ=amyloid-beta. *p<.05, FDR-corrected. **p≤.001, FDR-corrected. ^p<.05, uncorrected.

**Supplementary Table 11. Associations between tract integrity in cortical lobes and histopathological measures in amnestic and non-amnestic subtype.**

| | Cingulum | Frontal lobe | Temporal lobe | Parietal lobe | Occipital lobe | Insula |
|---|---|---|---|---|---|---|
| **MRI-histopathology association in amnestic AD** | | | | | | |
| **Tract FA** | | | | | | |
| | **Cingulum** | **Frontal lobe** | **Temporal lobe** | **Parietal lobe** | **Occipital lobe** | **Insula** |
| **ChAT cell density** | p=0.520 | p=0.628 | p=0.520 | p=0.520 | p=0.628 | p=0.628 |
| **Aβ load (%area)** | p=0.879 | p=0.330 | p=0.689 | p=0.879 | p=0.650 | p=0.650 |
| **p-tau load (%area)** | p=0.769 | p=0.769 | β= 0.107, p=0.018* | p=0.462 | p=0.769 | p=0.057 |
| **Tract MD** | | | | | | |
| | **Cingulum** | **Frontal lobe** | **Temporal lobe** | **Parietal lobe** | **Occipital lobe** | **Insula** |
| **ChAT cell density** | p=0.075 | β= -0.295, p=0.024* | β= -0.306, p=0.018* | β= -0.355, p=0.040* | p=0.241 | p=0.560 |
| **Aβ load (%area)** | p=0.977 | p=0.977 | p=0.977 | p=0.977 | p=0.977 | p=0.977 |
| **p-tau load (%area)** | p=0.923 | p=0.923 | p=0.923 | p=0.923 | p=0.923 | p=0.534 |
| **MRI-histopathology association in non-amnestic AD** | | | | | | |
| **Tract FA** | | | | | | |
| | **Cingulum** | **Frontal lobe** | **Temporal lobe** | **Parietal lobe** | **Occipital lobe** | **Insula** |
| **ChAT cell density** | p=0.786 | p=0.786 | p=0.839 | p=0.839 | p=0.786 | p=0.839 |
| **Aβ load (%area)** | p=0.844 | p=0.844 | p=0.844 | p=0.844 | p=0.844 | p=0.844 |
| **p-tau load (%area)** | p=0.827 | p=0.827 | p=0.827 | p=0.827 | p=0.827 | p=0.570 |
| **Tract MD** | | | | | | |
| | **Cingulum** | **Frontal lobe** | **Temporal lobe** | **Parietal lobe** | **Occipital lobe** | **Insula** |
| **ChAT cell density** | p=0.955 | p=0.955 | p=0.955 | p=0.955 | p=0.955 | p=0.955 |
| **Aβ load (%area)** | p=0.612 | p=0.634 | p=0.634 | p=0.612 | p=0.612 | p=0.634 |
| **p-tau load (%area)** | p=0.864 | p=0.864 | p=0.916 | p=0.864 | p=0.864 | p=0.864 |

Abbreviations: AD=Alzheimer's disease, FA=fractional anisotropy, MD=mean diffusivity, ChAT=cholinergic acetyltransferase, p-tau=phosphorylated-tau, Aβ=amyloid-beta. *$p<.05$, FDR-corrected.

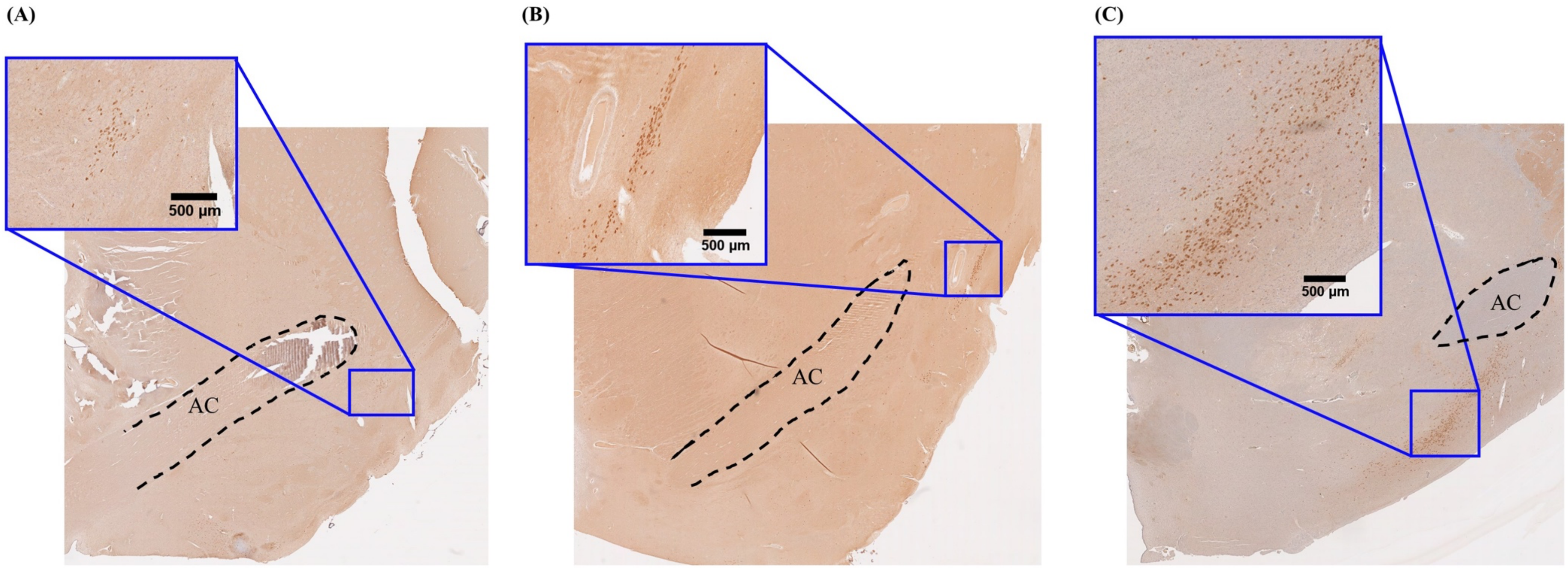


**Supplementary Figure 1. Subsectors of the NbM in three control donors.** The pre-anterior NbM of a control (78 year-old) showing ChAT cell reactivity at the lateral aspect and underneath the anterior commissure (AC, black dotted line) shown in (A). Anteroromediate NbM of a control (59 year-old) showing ChAT reactivity in a cluster at the lateral side of AC (black dotted line) shown in (B). Antero-intermediate NbM of a control (71 year-old) showing ChAT reactivity in a cluster underneath the AC (black dotted line) shown in (C). Abbreviations: µm=micron. AC=anterior commissure.

**Supplementary Figure 2. Selection of ChAT reactivity and pathology using automated ImageJ script.**

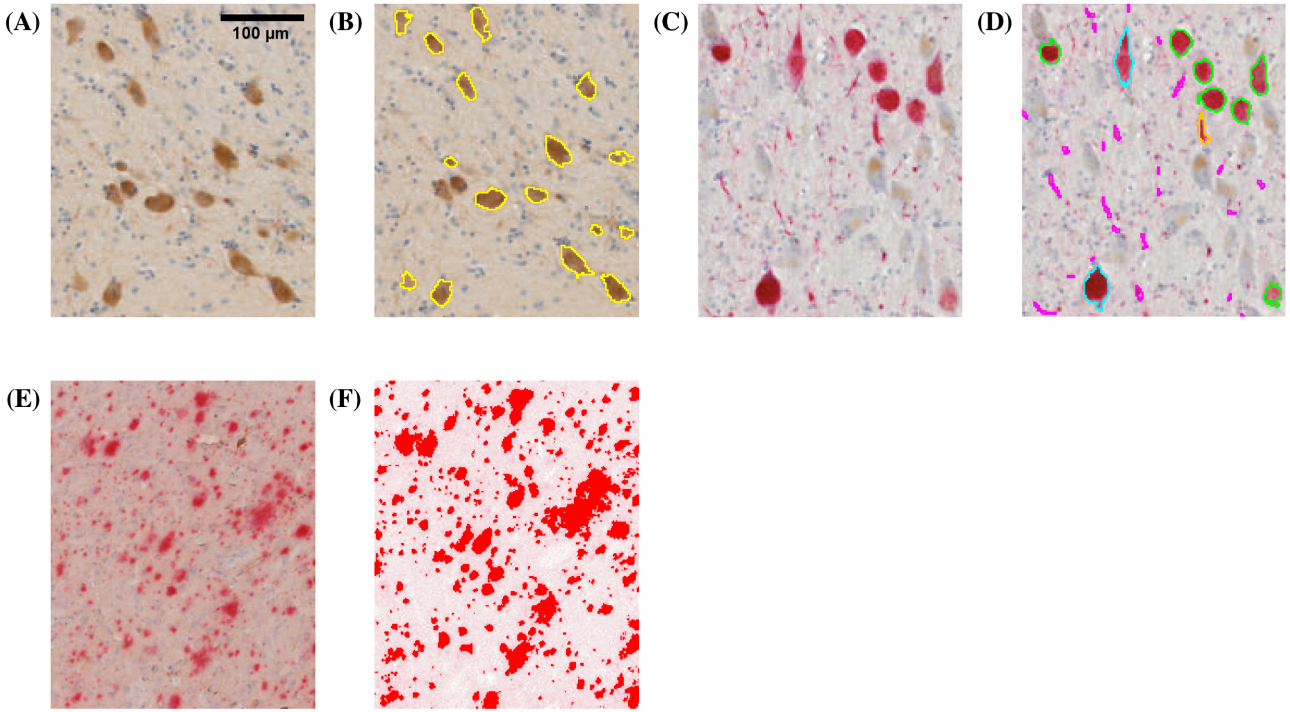


ChAT reactivity shown in brown ellipsoids in (A) were selected and counted for ChAT count, as shown in the yellow outlines in (B). p-tau positivity shown in red in (C) were selected in turquoise, green, orange and magenta as shown in (D) and further measured as percentage surface area for p-tau load. Aβ plaques positivity was measured as percentage surface area for Aβ load, as shown in (E) and (F). The scale bar in (A) applies to all images. Abbreviation: µm=micron.
